# Fast-spiking interneurons of the premotor cortex contribute to action planning

**DOI:** 10.1101/2021.05.14.443822

**Authors:** Nadia Giordano, Claudia Alia, Lorenzo Fruzzetti, Maria Pasquini, Alberto Mazzoni, Silvestro Micera, Leonardo Fogassi, Luca Bonini, Matteo Caleo

## Abstract

Planning and execution of voluntary movement depend on the contribution of distinct classes of neurons in primary motor and premotor areas. However, the specific functional role of GABAergic cells remains only partly understood. Here, electrophysiological and computational analyses are employed to compare directly the response properties of putative pyramidal (PNs) and fast-spiking, GABAergic neurons (FSNs) during licking and forelimb retraction in mice. Recordings from anterolateral motor cortex and rostral forelimb area, reveal that FSNs fire earlier and for a longer duration than PNs, with the exception of a subset of early-modulated PNs in deep layers. Computational analysis reveals that FSNs carry vastly more information than PNs about the onset of movement. While PNs differently modulate their discharge during distinct motor acts, most FSNs respond with a stereotyped increase in firing rate. Accordingly, the informational redundancy was greater among FSNs than PNs. These data suggest that a global rise of inhibition contributes to early action planning.

## INTRODUCTION

Preparatory activity occurring before the initiation of voluntary movements is critical for action planning and execution (Churchland, 2006; Godschalk et al., 1985; Guo et al., 2014b; Murakami et al., 2014; Weinrich and Wise, 1982; Wise and Mauritz, 1985). Specifically, premotor areas act as a conductor to orchestrate the network activity of the rest of the motor modules, on a moment-by-moment basis, and exhibit tuning for specific movements (Churchland, 2006; Churchland and Shenoy, 2007; Georgopoulos et al., 1982; Godschalk et al., 1985; Hocherman and Wise, 1991; Messier and Kalaska, 2000; Riehle and Requin, 1993). How do distinct neuronal classes contribute to this process? The anticipatory activity of pyramidal neurons (PNs) has been previously examined (reviewed in Svoboda and Li, 2018), however little is known on the contribution of GABAergic cells to these cortical computations (Isomura et al., 2009; Kaufman et al., 2013; Merchant et al., 2008).

Fast-spiking neurons (FSNs) are the most prevalent type of GABAergic interneurons in the cortex (Lourenço et al., 2020) and are well suited to shape neuronal dynamics (Isomura et al., 2009; Merchant et al., 2008; Pi et al., 2013; Polack et al., 2013; Sachidhanandam et al., 2016). FSNs exhibit thin action potentials and high spontaneous discharge rates (Merchant et al., 2012; Swanson and Maffei, 2019). In the rodent sensory cortex, FSNs contribute to sharpening the tuning of cortical neurons to preferred stimuli (Isaacson and Scanziani, 2011; Liu et al., 2011; Poo and Isaacson, 2009; Tan et al., 2011; Wu et al., 2008). In the primary motor cortex, they fire before PNs during spontaneous movements (Estebanez et al., 2017), suggesting a dynamic role of inhibition in shaping the tuning of PNs while routing information to the subsequent motor module (Merchant et al., 2008).

Here we recorded premotor neuronal activity from the anterior-lateral motor cortex (ALM, Chen et al., 2017; Guo et al., 2014b; Komiyama et al., 2010), which partially overlaps with the rostral forelimb area (RFA, Tennant et al., 2011; Vallone et al., 2016) in head-restrained mice allowed to either spontaneously lick or pull a handle in a robotic device (Spalletti et al., 2017). We found that PNs discharged later and for a shorter duration, except a small percentage of early modulated PNs in deep layers. PNs displayed specific activity during distinct motor acts, while most FSNs enhanced their discharge irrespective of the movement type. Computational analyses revealed that FSNs carried a greater amount of redundant information prior to PNs activation.

## RESULTS

### Activity of PNs and FSNs in the ALM during licking

To precisely target extracellular recordings within the ALM, we carried out initial optogenetic mapping experiments in 6 Thy1-ChR2 mice (Spalletti et al., 2017). We confirmed that the stimulation of ALM (data not shown), evoked mouth/tongue movements, as expected (Svoboda and Li, 2018).

Water-restricted, head-fixed mice were allowed to lick spontaneously drops of liquid reward, available through a drinking spout, centred in front of the animal, and triggering licking events. Offline, we categorized isolated, “single”, or “multiple” licks (**Figure 1A**, see Methods).

**Figure 1.**
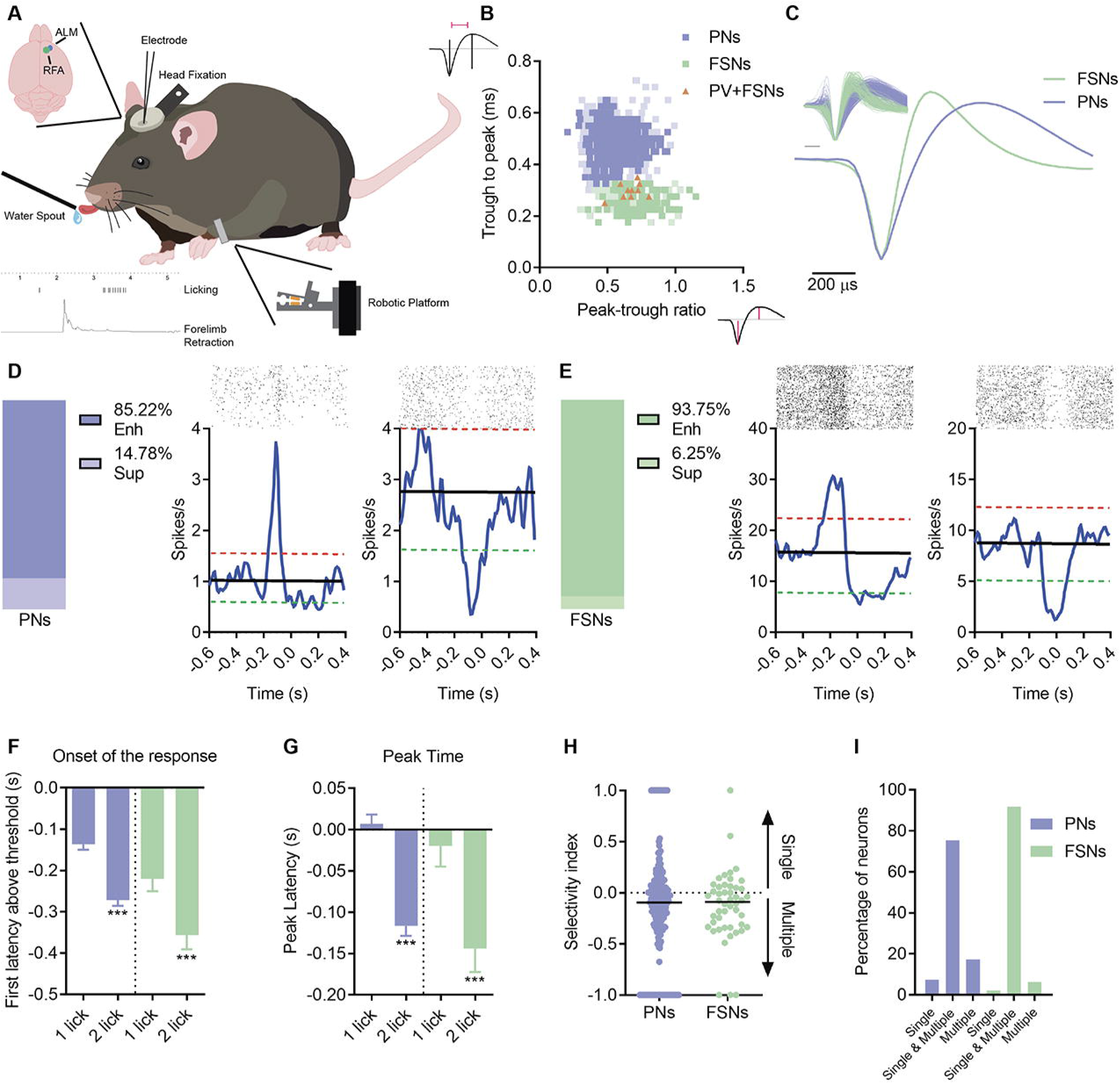
FSNs display less tuned responses during single vs multiple licks. (A) Schematic representation of a head-fixed mouse in the behavioral setup. In the bottom left, a scale bar of the licking behavior and a forelimb force peak are represented. (B) Scatter plot of spike waveform parameters for all units recorded (n = 1452). The violet and green filled squares represent individual putative PNs (PNs, task related or not, violet and light violet, respectively) and FSNs (FSNs, task related or not, green and light green, respectively). The orange filled triangles show spike shapes of individual PV+FSNs (activated at short latency by light). (C) Average spike waveforms for all units, PNs and FSNs, aligned to minimum and normalized by trough depth. All waveforms are displayed in the inset (top left). (D, E) Proportion of all responsive putative PNs – enhanced, violet, or suppressed, light violet – (D) and putative FSNs – enhanced, green, or suppressed, light green – during the licking activity (E). On the right, representative examples of raster plots and corresponding firing-rate-time-histograms showing enhanced (left) and suppressed (right) neurons. The red dotted lines represent the upper thresholds, the green dotted lines the lower ones, the black line is the mean baseline firing rate. Time = 0 corresponds to the first lick, not preceded by other licks for at least 0.6 s. PNs suppressed vs FSNs suppressed, Z-Test, z = 1.65, *p* = 0.09. (F, G) Histograms of the onset of the response (F) and the peak time (G) of PNs and FSNs obtained aligning PSTHs to the first or the second lick of a licking bout. Wilcoxon Test, \*\*\**p* < 0.001. (H) Selectivity of PNs and FSNs for single vs multiple licks. The peak of the response (spk/s) is used to calculate a SI (see Methods) ranging from −1 (neurons selective for multiple licks) to +1 (neurons selective for single licks). The black horizontal line corresponds to the median value. (I) Percentage of PNs and FSNs responsive to both single and multiple licks, or selective for single or multiple licks. Chi-square Test, *p* = 0.046. Single&Multiple PNs vs Single&Multiple FSNs, Z-Test, z = 2.5, *p* = 0.01.

We extracellularly recorded neuronal activity with either an acutely inserted single shank, 16-channels silicon probe (n = 10 mice, n = 693 units) or a chronic 16-microelectrodes array (n = 3 mice; n = 759 units,) from the ALM (and RFA). Spike detection and sorting were performed offline (Barthó et al., 2004; Mitchell et al., 2007; Niell and Stryker, 2010) to separate narrow- and broad-spiking neurons (**Figure 1B and 1C**).

Peristimulus time histograms (PSTHs) were created by aligning the spiking activity of each neuron to the first event of each licking bout (timed as “0”). For each PSTH, the mean firing rate was compared to thresholds (**Figure S1A**) to identify responsive neurons. Overall, in both acute and chronic recordings, we found n = 624 units (indicated by violet and green squares in **Figure 1B**) significantly modulated by movement (n = 828 PNs and FSNs not modulated by the task, indicated by light violet and light green squares, respectively; **Figure 1B** and **Table S1**). Responsive neurons with a thin spike shape displayed higher baseline discharge and shorter inter-spike intervals (ISI) than broad-spiking neurons (**Figures S1B and S1C**), consistent with their classification as putative FSNs. Broad-spiking neurons were instead considered as putative PNs.

To further validate the identification of FSNs, we performed extracellular recordings in mice expressing ChR2 selectively in Parvalbumin-positive, fast-spiking cells (Tantillo et al., 2020). FSNs waveforms were added to data for PNs/FSNs clustering. Notably, all the optogenetically tagged PV+FSNs displayed a small trough to peak time and peak-trough ratio, coherently with their identification as putative interneurons, thereby confirming the reliability of our identification method (**Figure 1B**).

In a first set of experiments (with acutely inserted silicon probes) in the ALM, we found that n = 251 neurons (203 putative PNs, 48 putative FSNs) showed a modulation of activity during licking behavior. The majority of putative PNs showed a firing rate enhanced by licking, and only 15% of them exhibited a decrease in firing rate during licking epochs (**Figure 1D**). The proportion of licking-suppressed FSNs tended to be lower (about 6%; **Figures 1E**). In the case of multiple licks, PNs and FSNs showed the maximum response modulation before the licking bout. This was clearly demonstrated by building mean PSTHs aligned with the first and the second lick in the series. The alignment with the second lick showed that both onset of the response and peak latency were shifted about 0.150 s earlier than after alignment with the first lick (**Figures 1F and 1G**). These data suggest that neuronal discharges of both PNs and FSNs in the ALM are mainly related to the planning of entire licking bouts rather than the execution of each lick of the series.

We next compared the discharges of PNs and FSNs during either single or multiple licks. We computed a selectivity index (SI), ranging from −1 to 1, based on the peak of neuronal modulation, indicating the degree of tuning of each neuron for different licking strategies (**Figure 1H**, 1 indicates exclusive selectivity for single licks, −1 for multiple licks). We then plotted the percentages of PNs and FSNs modulated only during single licks (SI = 1), multiple licks (SI = −1), or during both (−1 < SI < 1; **Figure 1I**). The statistical analysis revealed that the distribution was different for PNs vs FSNs, and the proportion of neurons responsive for both single and multiple licks was greater for FSNs. These data indicate a more specific activity of PNs compared to FSNs, which showed instead a broader tuning.

### FSNs show earlier and more sustained activation than PNs during licking

We next investigated PNs and FSNs firing activity during single and multiple licks. First, we analyzed the onset of the (enhanced or suppressed) response (see **Figure S1A**), finding for the majority of recorded neurons a positive or negative variation prior to movement onset, independently from the forthcoming licking strategy (**Figure 2A**). Onset of PN discharge was earlier in multiple than in single licks (**Figure 2A**). FSNs fired ~ 0.1 s earlier than PNs, similarly across the two behavioral conditions. A cumulative distribution curve of the onset for individual neurons demonstrated a clear shift to the left for FSNs vs PNs (**Figures 2B and 2C**), indicative of an earlier recruitment of FSNs.

**Figure 2.**
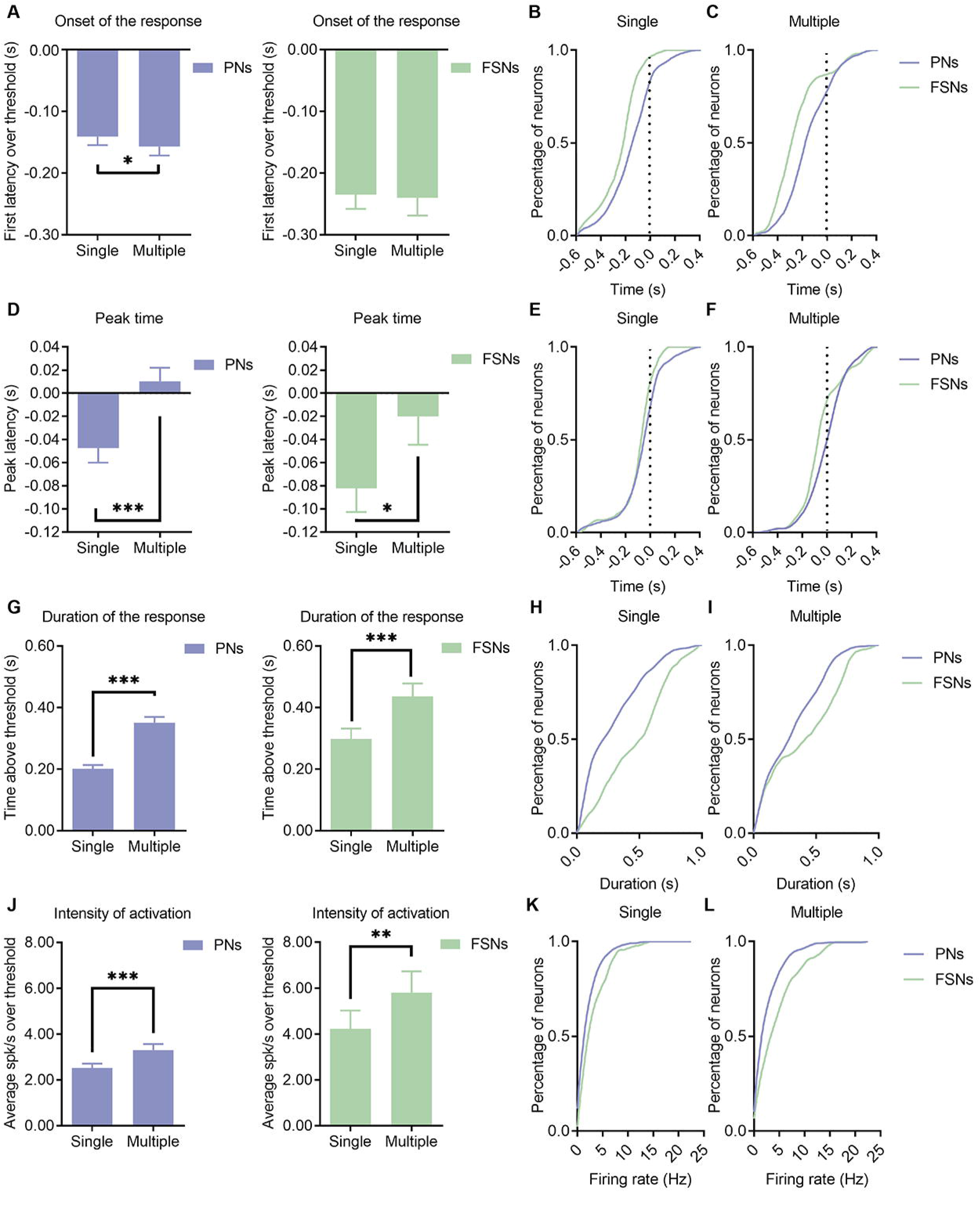
FSNs show earlier and more sustained activation than PNs during licking. (A) Histograms of onset of the response, defined as the first latency above or below the thresholds on PSTHs, for PNs (left) and FSNs (right), during single and multiple licks. Wilcoxon Test, \**p* < 0.05. (B, C) Cumulative distribution of the onset of the response for all PNs and FSNs during a single isolated lick (B) or multiple licks (C). K-S Test, Single, *p* = 0.001, Multiple, *p* < 0.001. (D) Histograms of the peak time of PNs (left) and FSNs (right) during single and multiple licks. The peak discharge is significantly delayed for both PNs and FSNs during multiple licks. Wilcoxon Test, *p* < 0.05, \*\*\**p* < 0.001. (E, F) Cumulative distribution of the peak time for all PNs and FSNs during a single isolated lick (E) or consecutive multiple licks (F). K-S Test, Single, *p* = 0.064, Multiple, *p* = 0.0063. (G) Histograms of the duration of the response of PNs (left) and FSNs (right), during single and multiple licks. Wilcoxon Test, \*\*\**p* < 0.001. (H, I) Cumulative distribution of the duration of the response for all PNs and FSNs during a single isolated lick (H) and consecutive multiple licks (I). K-S Test, Single, *p* = 0.0158, Multiple, *p* = 0. 0269. (J) Histograms of the intensity of activity of PNs (left) and FSNs (right), during single and multiple licks. Wilcoxon Test, PNs, \*\**p* < 0.01, \*\*\**p* < 0.001. (K, L) Cumulative distribution of the intensity of activation for all PNs and FSNs during a single isolated lick (K) and multiple licks (L). K-S Test, Single, *p* = 0.065, Multiple, *p* = 0.0051.

Then, we examined the timing of the peak of activity (or suppression) for each neuron. In multiple licks, the average peak time was delayed for both PNs and FSNs (**Figure 2D**). Cumulative distributions of the peak latency are reported in **Figures 2E and 2F**. A robust statistical difference between PNs and FSNs was present for multiple licks (**Figure 2F**). Specifically, half of PNs reached their maximum firing rate before the onset of the licking bout, whereas about 75% of FSNs had their peak of activity prior to licking onset (**Figure 2F**).

Next, we explored the duration of neuronal response, greater for both PNs and FSNs when mice performed multiple vs single licks (**Figure 2G**). Interestingly, the response duration was longer in FSNs during both single and multiple licks as compared to PNs (**Figures 2H and 2I**).

Similar results were obtained by analyzing the magnitude of the activation of the two neuronal classes. During multiple licks, both PNs and FSNs showed greater discharge than during a single lick (**Figure 2J**). Furthermore, the FSNs displayed a higher activity relative to PNs, and this was more evident in multiple than in single licks (**Figures 2K and 2L**).

Altogether, these findings suggest the idea that single or multiple licks are coded by the differential activity patterns of both PNs and FSNs in terms of onset, peak discharge, duration and magnitude of neuronal activity.

### Information content of firing rate

The distinct behavior of PNs and FSNs reported in the previous section can be clearly appreciated by building mean PSTHs for the two classes of neurons (**Figures 3A and 3B**).

**Figure 3.**
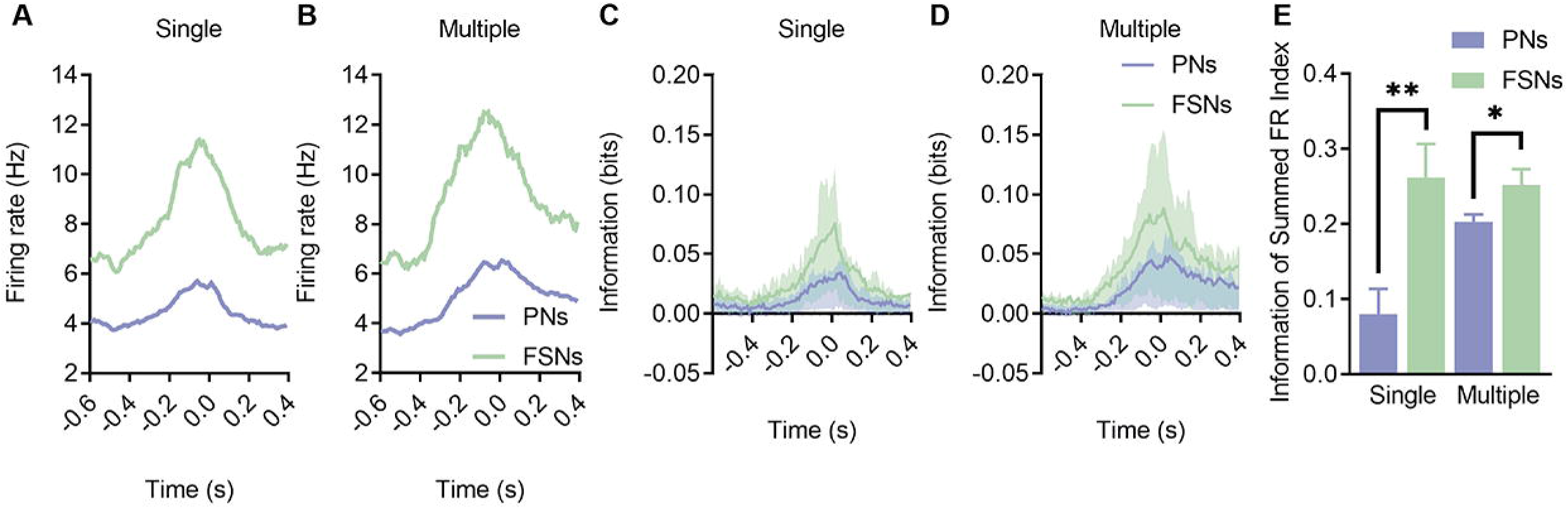
FSNs convey a considerable amount of information and prior to PNs activation. (A, B) Average PSTHs for all PNs (violet) and FSNs (green) in a 1 s window (0.6 s before and 0.4 s after the licking event) during single (A) and multiple (B) licks. (C, D) Information carried by firing rate of PNs (violet) and FSNs (green) about the presence of single (C) and multiple (D) licks. Information is computed over 0.05 s bins (with a sliding time window of 0.01 s width) in a 1 s window (0.6 s before and 0.4 s after the licking event). Lower and higher shades represent, respectively, the 25 and 75 percentile. Wilcoxon Test, *p* < 0.001. (E) Information of summed firing rate index for couple of PNs and FSNs of the same recording session for both single and multiple licks. Mann-Whitney Test, *\*p* < 0.05, \*\**p* < 0.01.

We next computed, for all the recorded units, the mutual information between the firing rate and the behavioral states (i.e., rest, single lick and multiple licks; see Methods). The fraction of informative neurons was 0.74 for FSNs and 0.63 for PNs. Within the subset of informative neurons, FSNs carried vastly more information than PNs about the onset of both single (0.074 bits, PNs, 0.130 bits, FSNs) and multiple licks (0.140 bits, PNs, 0.221 bits, FSNs).

Coherently with an earlier onset of the response, FSNs information content ramped up earlier than PNs (**Figures 3C and 3D**). Information carried by FSNs became 3 SD larger than baseline for at least two consecutive bins ~0.05 s earlier than PNs. Comparing single lick and rest, the information exceeded the threshold 0.25 s before movement onset in FSNs and 0.2 s in PNs. The peak was reached at licking onset in FSNs and 0.03 s later in PNs. Multiple licks vs rest yielded similar results: the information exceeded the threshold 0.33 s before the onset of the movement in FSNs and 0.27 s in PNs. Peak was reached 0.02 s in FSNs and 0.05 s in PNs after the event. These dynamics were similar to the responsivity illustrated by the PSTHs results (compare **Figures 3C and 3D** with **Figures 3A and 3B**).

We then computed animal-wise the amount of information carried by the summed firing rate of the recorded FSNs and PNs population and found that FSNs carried more redundant information. The Information of summed firing rate index (see Methods) is significantly higher for FSNs than for PNs (mean 0.08 for PNs, 0.26 for FSNs, single lick vs rest; mean 0.20 for PNs, 0.25 for FSNs, multiple licks vs rest; **Figure 3E**).

Overall, these results suggest that the local firing rate of FSNs conveys a considerable amount of information prior to PNs activation, further supporting the idea that a robust and coherent inhibitory activity might be important during the planning of the movement.

### Layer-specific responses of PNs and FSNs

Linear probes allowed us to investigate the laminar distribution of recorded neurons. Specifically, units were classified as superficial (channels 1-8, ~ < 600 μm depth) or deep (channels 9-16, ~ > 600 μm depth) neurons. In our sample, about 25% of PNs and FSNs were recorded from superficial layers. **Figures S2A and S2B** report the onset of activity for each recorded unit as a function of depth (i.e. channel number). While the average response onset of FSNs precedes the one of PNs (consistent with **Figures 2A–2C**), a small proportion of PNs appear to increase their firing rate earlier, simultaneously with FSNs (**Figures S2A and S2B**).

By arbitrarily splitting the temporal window before licking onset into two identical segments of 0.3 s each (−0.6/−0.3 s and −0.3/0 s), we found that early-birds PNs appeared to be more prevalent in deep than in superficial layers (**Figures S2C and S2D**).

These results confirm that preparatory activity begins in deep layers of ALM (Chen et al., 2017), and involves both FSNs and PNs.

### Direct comparison of the neuronal responses of PNs and FSNs during two motor acts

Early and sustained inhibition by FSNs during licking preparation may be a general mechanism that contributes to action selection during the preparatory phase prior to movement onset, regardless of the effector to be used for acting. To test this hypothesis, we compared the activity of the same FSNs and PNs during two types of motor tasks, i.e. licking and forelimb retraction. We took advantage of a robotic platform (M-Platform, Allegra Mascaro et al., 2019; Spalletti et al., 2017), which allows mice to perform several trials of forelimb pulling. Neuronal discharges were aligned on the onset of force peaks (Pasquini et al., 2018; Spalletti et al., 2014). Animals also performed spontaneous licking within the same experimental session. In the following sections, we describe the neuronal discharges during “isolated” (i.e. spaced by more than 1 s) pulling and multiple licking events.

For these experiments we employed a chronic array, centered on the ALM but exceeding the boundary with the adjacent RFA (Alia et al., 2016; Tennant et al., 2011). Electrode contacts were positioned in deep layers; we isolated n = 373 units (PNs, n = 313; FSNs, n = 60; mice, n = 3), which were responsive to either licking or pulling (or both).

We found a higher proportion of neurons whose discharge was suppressed by movement, with respect to previous data collected in acute recordings. Specifically, 40% of PNs, whose discharge was modulated during forelimb retraction, showed movement-related suppression of their discharge; a similar proportion (37.0%) of PNs responsive for licking behavior were also suppressed. For FSNs, the percentages of suppressed neurons were similar (39.1%) for forelimb retraction, and lower (20.3%) for licking. These data suggest that corticofugal pyramidal neurons as well as FSNs located in deep layers are particularly susceptible to movement-related suppression. Therefore, we analyzed enhanced and suppressed neurons separately (**Table S2**).

### FSNs exhibit lower selectivity than PNs for licking behavior and forelimb retraction

Neuronal selectivity for each type of movement (i.e., licking vs pulling) was quantified based on the amplitude of the discharge modulation by means of a SI ranging from −1 (pure selectivity for forelimb retraction) to 1 (pure selectivity for licking; **Figure 4A**). Licking was the preferred neuronal response for all recorded units, consistent with the position of the array. We categorized neurons into either “forelimb-specific”, “licking-specific”, and “forelimb+licking” based on their SI. Overall, we found that PNs displayed more variability in their responses to the two different motor acts compared to FSNs, that were less tuned than PNs (**Figure 4B**), consistent with the data previously reported for “single” and “multiple” licks (**Figure 1I**).

**Figure 4.**
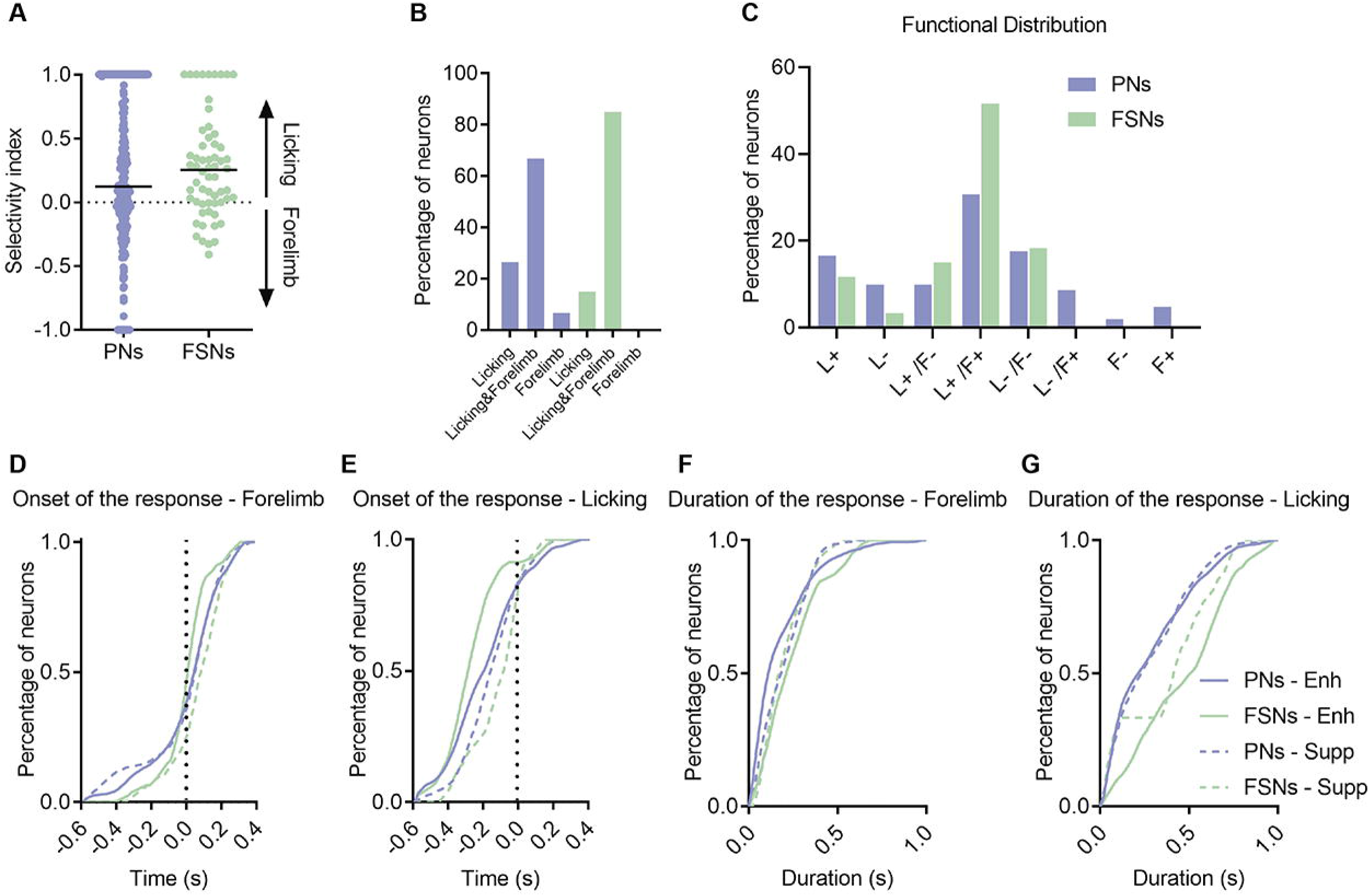
FSNs exhibit lower selectivity than PNs for licking behavior and forelimb retraction. (A) The peak of the response (spk/s) is used to calculate a SI ranging from −1 (neurons selective for forelimb retraction) to +1 (neurons selective for licking behavior). The black horizontal line corresponds to the median value. (B) Percentage of PNs and FSNs responsive to both licking and forelimb retraction, or selective for licking or forelimb pulling. Chi-square test, *p* = 0.01. (C) Functional distribution of neurons responsive for licking (L), forelimb pulling (F) or both of them (L/F), classified as enhanced (+) or suppressed (-) by the movement. Chi-square test, *p* = 0.0052. (D) Cumulative distribution of the onset of the response for all neurons during the forelimb retraction. Enhanced PNs vs suppressed PNs, K-S Test, *p* = 0.91. Enhanced FSNs vs suppressed FSNs, K-S Test, *p* = 0.12. Enhanced PNs vs enhanced FSNs, K-S Test, *p* = 0.081. (E) Cumulative distribution of the onset of the response for all neurons during a licking bout. Enhanced PNs vs suppressed PNs, K-S Test, *p* = 0.043. Enhanced FSNs vs suppressed FSNs, K-S Test, *p* = 0.0090. Enhanced PNs vs enhanced FSNs, K-S Test, p = 0.0069. (F) Cumulative distribution of the duration of the response for all neurons during the forelimb retraction. Enhanced PNs vs enhanced FSNs, K-S Test, *p* = 0.029. Enhanced FSNs vs suppressed FSNs, K-S Test, *p* = 0.216. Enhanced PNs vs suppressed PNs, K-S Test, *p* = 0.137. (G) Cumulative distribution of the duration of the response for all neurons during a licking bout. Enhanced PNs vs enhanced FSNs, K-S Test, *p* = 0.0009. Enhanced FSNs vs suppressed FSNs, K-S Test, *p* = 0.610. Enhanced PNs vs suppressed PNs, K-S Test, *p* = 0.987. Enhanced neurons are represented as continuous lines (PNs, violet; FSN, green), dotted lines indicate the suppressed PNs and FSNs.

To further investigate the response properties of PNs and FSNs during the two motor tasks, we subdivided recorded units into different functional classes, according to the movement-induced modulation of their discharge. Specifically, neurons responsive to only one type of movement were classified as enhanced/suppressed by licking (L+, L-) or forelimb pulling (F+, F-). Neurons responsive to both movements showed either a mutual (L+/F+, L-/F-) or opposite modulation (L+/F-, L-/F+) during each motor task. We found that PNs (violet bars in **Figure 4C**) were distributed across all functional classes. In contrast, the vast majority of FSNs (> 72%) were mutually modulated (i.e., suppressed or enhanced) by the two different movements (i.e., L+/F+, 50% and L-/F-, 20%). The functional distributions of PNs and FSNs were significantly different (**Figure 4C**).

We next compared response onset and duration among the different populations of neurons. Enhanced FSNs started to discharge before facilitated PNs (**Figures 4D and 4E**). Note, however, that, consistently with the laminar recordings (**Figure S2**), a small subset of pyramidal neurons (approx. 15%) modulated their discharges very early, especially during forelimb retraction (**Figure 4D**). Interestingly, the suppressed FSNs showed a delayed discharge onset relative to the enhanced FSNs, especially during licking (**Figures 4D and 4E**).

In terms of duration of the response, this was significantly greater for the FSNs, specifically those excited by movement, considering both pulling (**Figure 4F**) and licking (**Figure 4G**). The suppressed FSNs showed a shorter duration of modulation, although not statistically different from that of enhanced FSNs (**Figures 4F and 4G**). There was no difference in the discharge duration between enhanced and suppressed PNs (**Figures 4F and 4G**).

Altogether, these data concur with the previous laminar recordings indicating an early and prolonged discharge of FSNs activated by either licking or pulling. Interestingly, the suppressed FSNs were modulated at longer latencies during movement generation.

## DISCUSSION

In summary, our data reveal that FSNs fire in anticipation of PNs within the same cortical module. Specifically, they start to modulate and reach their peak of activity earlier than FSNs. These findings are in agreement with a previous electrophysiological study examining discharges of FSNs and PNs in mouse primary motor cortex during sensory-triggered as well as voluntary forelimb reaching (Estebanez et al., 2017). Thus, the early involvement of FSNs appears to be a hallmark of both primary motor and premotor areas. Interneurons also appear to increase their firing rates more than putative PNs during movement planning and execution. Duration of activity is also greater for FSNs as compared to PNs. Overall, FSN firing rates appear to carry more information about movement onset than PNs. Preparatory/ramping activity in ALM PNs has been shown to be maintained by a recurrent excitatory loop that involves both the cortex and the ipsilateral thalamus (Guo et al., 2017). Since FSNs are directly reached by thalamic afferents (Lourenço et al., 2020), this recurrent thalamocortical loop may sustain persistent firing activity in FSNs.

It is worth noting that, although PNs were recruited later than FSNs during motor planning, a fraction of them located in deep layers was early-modulated. These data suggest that such PNs may represent preparatory “master” neurons that subsequently command downstream, more executive PNs and FSNs. In keeping with our results, it has been shown that preparatory activity appears first in deep layers of ALM (Chen et al., 2017). For what concerns FSNs suppressed by movement, the onset data clearly show that they are consistently delayed with respect to the other populations. Since PV+FSNs form a highly interconnected set of neurons (Lourenço et al., 2020), it is likely that the suppressed fast-spiking population receive direct synaptic input from enhanced FSNs.

PNs and FSNs recorded exhibited robust differences in tuning for the type of movement. FSNs were less selective for movement type than PNs (i.e. multiple vs single licks), while PNs showed a variety of behaviors and were distributed in several functional classes, with enhancement or suppression of discharge depending on the specific movement (i.e. licking and pulling). On the contrary, the percentage of suppressed FSNs was lower, and they often increased their firing rate during both pulling and licking. Accordingly, FSNs appear to carry more redundant information than PNs, consistently with the fact that FSNs are synchronized by electrical and chemical synapses (Lourenco et al., 2020). In the prefrontal cortex of mice performing a sensory discrimination task, PV+FSNs are activated by all task-related events (sensory cues, motor action, and trial outcomes), while responses of PNs are diverse and more selective (Pinto and Dan, 2015). The broader tuning of FSNs is consistent with data in sensory cortices, where interneurons are poorly selective for stimulus features, such as orientation selectivity (Hofer et al., 2011; Kerlin et al., 2010), as well as in the monkey parieto-premotor cortices, as shown by recent evidence concerning visual and motor tuning for object type during visually-guided grasping actions (Ferroni et al. 2021, *in press, Current Biology).*

It has been hypothesized that the activity of interneurons, including FSNs, provides an inhibitory gate that prevents preparatory activity from causing undesired movements. If this were the case, interneuron firing rates should be reduced around the time of movement, which was not observed in the present experiments. Another possibility is that FSN-mediated inhibition may serve to suppress other actions (e.g., movement of other body parts). If FSNs act to prevent adjacent cortical modules from producing other movements, one would predict the existence of distinct licking- and forelimb-related FSNs which reciprocally inhibit the respective PNs. However, our data do not provide support for such a model, as more than 50% of FSNs increase their discharge during both licking and forelimb retraction. Thus, a sustained, overall rise in FSN activity appears to be required during action planning, likely to reach a critical level of inhibition for proper execution of the subsequent lick/forelimb pulling.

Several hypotheses may be put forward to explain the early, prolonged and broadly tuned activation of FSNs reported in the present study. (i) The discharge of FSNs may be required to sculpt the response selectivity of nearby PNs. In the motor cortex, the magnitude of inhibition directly affects tuning of individual PNs before and during movement execution both in mice (Galiñanes et al., 2018) and nonhuman primates (Georgopoulos et al., 1982; Merchant et al., 2008). (ii) Activity of FSNs might provide an inhibitory constraint that maintains firing rates of PNs within an “optimal subspace” (Afshar et al., 2011) that allows accurate movement (Churchland, 2006). Experimental testing of these and other possibilities will require optogenetic modulation of the activity of FSNs at specific times of motor planning in delayed response tasks (Svoboda and Li, 2018).

Altogether, the present data reveal an early and prolonged involvement of FSNs during movement planning in premotor areas, which may play a role in sculpting response selectivity of nearby pyramidal neurons. Future studies leveraging causal techniques should elucidate the key role of inhibition in shaping PN response properties.

## Acknowledgements

We thank Francesca Biondi (CNR PI) for excellent animal care. This project has received funding from the H2020 EXCELLENT SCIENCE-European Research Council (ERC) under grant agreement 692943 (BrainBIT). This work was also funded by Regione Toscana (PERSONA project, bando Salute 2018) and by Fondazione Cassa di Risparmio di Padova e Rovigo (Foundation Cariparo, project #52000 to M.C.).

## Author contributions

Conceptualization, NG, CA and MC; Methodology, NG and CA; Investigation, NG and CA; Software LFr and MP; Formal Analysis NG, CA and LFr; Writing – Original Draft, NG, CA and MC; Writing – Review & Editing, NG, CA, LB, LFo, and MC; Funding Acquisition, MC; Supervision, AM, SM, LB and LFo

## METHODS

### EXPERIMENTAL MODELS AND SUBJECT DETAILS

All experiments were carried out in accordance with the EU Council Directive 2010/63/EU on the protection of animals used for scientific purposes, and were approved by the Italian Ministry of Health (authorization number 684/2020-PR). Animals were housed in rooms at 22 °C with a standard 12h light/dark cycle. Food (standard diet, 4RF25 GLP Certificate, Mucedola) and water were available ad libitum, except for the experimental period, during which mice were water deprived overnight. Electrophysiological recordings were conducted on 13 subjects. Experiments were carried out on 3–5 months old wild-type (C57BL6/J) male mice. For optotagging of FSNs, 2 PV-Cre mice (Tanahira et al., 2009) (B6;129P2-Pvalb tm1(cre)Arbr/J, Jackson Laboratories, USA) were injected with 600 nl of an AAV vector (AAV1 .EF1 .dflox.hChR2(H134R)-mCherry.WPRE.hGH, Addgene, USA), into the motor cortex. The AAV vector contains the doublefloxed channelrhodopsin-2 (ChR2) gene, which is thus expressed in parvalbumin interneurons through Cre-mediated recombination (Spalletti et al., 2017; Tantillo et al., 2020). We referred to them as PV+FSNs (**Figure 1B**). Six B6.Cg-Tg(Thy1-COP4/EYFP)18Gfng (ChR2) mice expressing ChR2 mainly in corticofugal, layer V neurons were used to map mouth/tongue movements in the ALM.

#### Surgery procedure and animal preparation

Mice were deeply anesthetized with an intraperitoneal injection of avertin (0.020 ml/g), and positioned on a stereotaxic frame; the scalp was partially removed, the skull cleaned and dried. A ground screw was implanted above the cerebellum. Mice were implanted with a custom-made lightweight head post, placed on the skull on the left hemisphere, aligned with the sagittal suture and cemented in place with a dental adhesive system (Super-Bond C&B). A thin layer of the dental cement was used to cover the entire exposed skull.

For acute recordings, a recording chamber was built using a dental cement (resin adhesive cement, Ivoclar Vivadent) and centered on the right ALM (1.8□ mm lateral and 2.5□mm anterior to Bregma).

The skull over the recording area was covered by sterile low melting agarose Type III (A6138, Sigma-Aldrich, Inc.) and sealed with Kwik-Cast (WPI). On the day before the first acute recording session, 6 B6.Cg-Tg(Thy1-COP4/EYFP)18Gfng (ChR2) mice were anaesthetized with ketamine (100 mg/kg) and xylazine (10 mg/kg) cocktail, the chamber covering removed and cortex was optogenetically stimulated following a grid with nodes spaced 500 μm. For each site, optogenetic stimulation (3 ms single pulses, 0.2Hz) was delivered by means of PlexBright Optogenetic Stimulation System (PlexonInc, USA) with a PlexBright LD-1 Single Channel LED Driver (PlexonInc, USA) and a 473 nm Table-top LED Module connected to a 200 μm Core 0.39 NA optic fiber (ThorlabsInc, USA). Movements of tongue/mouth were collected by a second experimenter, blinded to the stimulation coordinates. A small craniotomy (diameter, 0.5 mm) was then performed over sites where the larger tongue/mouth movements could be evoked. In wild-type mice, the craniotomy was performed in the same region of Thy1-ChR2 mice. Finally, the chamber was filled with agarose and sealed.

For chronic implants, a craniotomy (diameter, 0.8 mm) was made over the right ALM, partially covering the Rostral Forelimb Area (RFA, 1.2 mm lateral and 2 mm anterior to bregma). A planar multi shank 4×4 array (Microprobes for Life Science) was positioned over the craniotomy and microwires were inserted into the cortex, up to ~800 μm depth. Then the craniotomy was covered with low melting agarose and the array fixed and embedded with dental cement (Super-Bond C&B and Paladur). Mice were allowed to awaken and then housed separately.

#### Behavioral training and data analysis

After recovering from surgery, mice were water restricted in their home cages, with food still available. Condensed milk was provided as a reward during the tasks and mice were also provided with water ad libitum for about 1 hour/day, following each recording session.

During the shaping phase, mice were placed in a U-shaped restrainer (3 cm inner diameter), head-fixed through the metal post cemented on their skull and habituated to lick drops provided randomly by the experimenter through a feeding needle mounted on a piezoelectric sensor sensing the movement of the tongue.

Each shaping session lasted from 15 min up to 60 min for at least 3 consecutive days. Digital signals from the licksensor provided information about the licking movements directly to the recording apparatus. Licking events were classified as either single or multiple licks. The start lick was defined as a movement that was not preceded (for at least 0.6 s) by any other licking event. A single lick was a start lick not followed by any other lick for at least 0.6 s; multiple licks were defined as start licks followed by at least two other consecutive licks (≤ 0.4 s among consecutive licks). Time intervals lasting for ≥ 1 s and distant at least 0.5 s from the end or the start of licking trials were considered as resting intervals and used as a baseline for the analysis of neural activity.

For identification of PV neurons in PV-cre mice, the site of AAV injection was illuminated with an optic fiber (200μm Core 0.39 NA, Thorlabs, USA). Optogenetic stimulation (50 0.2 s pulses, 0.2 Hz) was delivered by means of PlexBright System (Plexon, USA) with a PlexBright LD-1 Single Channel LED Driver (Plexon, USA) and a 473 nm Table-top LED Module. After spike sorting, PV-positive neurons were defined as neurons increasing their firing rate by 5 ms from the beginning of the blue light pulse (i.e. ChR2-positive neurons) and with a sustained activity for the entire stimulation length (i.e. FSNs).

For the chronic recordings, in which forelimb-driven response was also assessed, mice were shaped and trained on a robotic platform, the M-Platform (Spalletti et al., 2014) that comprises a 1-axis load-cell, a linear slide connected to a custom-designed handle that was fastened to the left wrist of the animal. During recording sessions, the forepaw, contralateral to the implanted ALM, was maintained in a slightly extended position and the force peaks exerted to attempt retractions were detected by the load-cell and offline aligned with neural signals.

#### Single-unit recording and spike sorting

The electrophysiological data were continuously sampled at 40 kHz and bandpass filtered (300 Hz to 6 kHz), using a 16-channel Omniplex recording system (Plexon, Dallas, TX).

For acute recordings, a NeuroNexus Technologies 16-channel linear silicon probe with a singleshank (A1×16-3mm-50-177, 50μm spacing among contacts) was slowly lowered into the ALM; the tip of the probe was placed at about 1000 μm depth using a fine micromanipulator (IVM, Scientifica). The recording chamber was filled with sterile saline solution (NaCl 0.9%). Before the beginning of the recording, the electrode was allowed to settle for about 10 min. For each animal, a number of one up to seven extracellular recording sessions were performed.

For chronic recordings, mice were recorded on up to 10-15 recording daily sessions per animal over a 15 days period.

The extracellular recording data were processed to isolate spike events by a spike sorting software (Offline Sorter™ v3.3.5, Plexon), using principal component analysis; events (spike-detection interval > 1.0 ms) that exceeded a 4 SDs threshold above the background were sorted. The spike waveforms were aligned at global minimum and the artifact waveforms were removed. The single-unit clusters were manually defined.

#### Data analysis

The recorded units were classified based on their average waveforms into putative pyramidal neurons (PNs) and putative fast-spiking neurons (FSNs). Two waveform parameters were used for the classification: the ratio between the height of the maximum peak and the initial negative trough, and the trough-peak time. A k-means clustering was applied. The clustering was verified by optogenetic tagging of PV-positive neurons.

The relation between single neuron activity and the events of the behavioral task was analysed using MATLAB (MathWorks). Peristimulus Time Histograms (PSTHs) were built aligning spike events on the start lick, for both single and multiple licks, and on the onset of the force during forelimb pulling. Only intervals with stable unit activity were included and spikes were averaged over 0.05 s with 0.01 s steps. The PSTH covered a time window of 1 s, from 0.6 s before the starting event (lick or force onset) and 0.4 s after it. Responsive neurons were identified by comparing firing activity in the PSTHs with the mean firing rate and an upper and lower threshold, calculated during resting periods (lasting ≥ 1 s, and distant from event trials ≥ 0.5 s). Bootstrapping was used to estimate the thresholds; lower and upper thresholds were, respectively, the 2.5 and 97.5 percentile of the probability distribution function obtained during the resting intervals. A unit was considered responsive for the licking behaviour or forelimb retraction when, for at least three consecutive bins (0.03 s), its firing rate went over (enhanced neurons) or under (suppressed neurons) the considered thresholds.

The onset of activity was defined as the first bin of the ≥ 3 consecutive bins above/below the upper/lower threshold; the time of the bin in which the firing rate (spk/s) was maximum/minimum was considered as the peak time. Selectivity indices were measured considering the peak firing rate (spk/s); they were defined as:

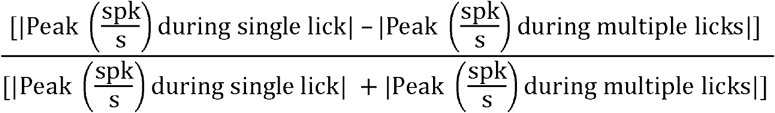

and:

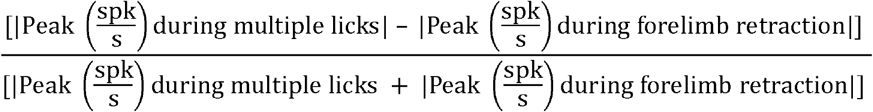

The duration of the response was the number of bins above/below the upper/lower threshold. The intensity of activation was defined as:

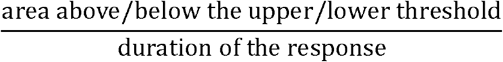

#### Information content

We measured the information content (Shannon, 1948), carried out by the mean firing rate of each neuron about two different sets of conditions. Set 1: 0.8 s intervals centred at single licks (see above) vs rest, i.e. randomly selected 0.8 s intervals during which animals were at rest, distant at least 1.5 s from other licking or rest intervals. Set 2: 0.8 s intervals centred at the onset of multiple licks (see above) vs rest.

The mean firing rate (`mfr`) associated to each trial was measured over the whole window. The mutual information of summed firing rates (`E, Mfr`) between `mfr` and each set of events E was computed as follows:

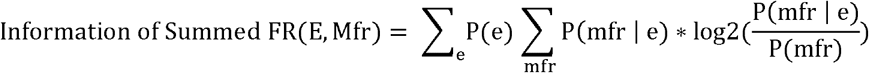

Where P(e) was the probability of the presentation of the specific event e, P(m) the probability over all trials and all conditions of the neuron to have the mean firing rate mfr in a given interval, `P(mfr | e)` the probability of the mean firing rate `mfr` to be associated to the event e. Mean firing rates were binned in N equipopulated bins, where N was the minimum value between the square root of the number total number of trials and the number of unique values in the array of mean firing rates.

To reduce the bias in the estimation of the information due to the limited dataset, a quadratic extrapolation method was used (Panzeri et al., 2007). A statistically significant threshold was obtained bootstrapping 100 times (shuffling the conditions associated to each trial), and, for a major solidity, only neurons with an IC > 95 percentiles of the bootstrapped distribution, in at least one of the two combinations, were included, generating a subset of informative neurons.

We also calculated the information content over time: we considered 0.8 s before and 0.4 s after the licking event, and we computed a local mean firing rate (LMF) over a moving average of 50 ms with steps of 10 ms. Then, for each step we repeated the procedure described above. For this analysis we only used the subset of informative neurons described above.

For each recording session, we computed animal-wise the amount of information carried by summed firing rates of the recorded FSNs and PNs population. Each recording session has a different number of neurons and a different ratio between FSNs and PNs, for this reason, to be able to compare results from different recording sessions, the information of summed firing rates was computed considering *N* couples of neurons belonging to the same class for each recording. *N* was the minimum value between all the possible combinations of same-class-neurons and 40.

For each couple of neurons, information of summed firing rates was calculated with the following equation:

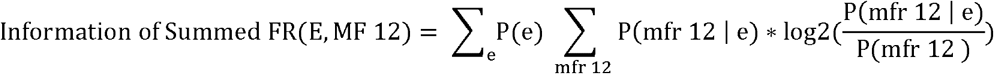

Where `Information of Summed FR(E, ISF 12)` is the information given by the summed firing rates of neuron 1 and 2, `P(e)` was the probability of the presentation of the specific event e, `P(mfr 12)` the probability that the sum firing rate of the neurons to have the mean firing rate `mfr 12` over all trials of all conditions, `P(mfr 12 | e)` is the probability of the mean firing rate `mfr 12` to occur during the event `e`.

We used the same bias correction method and the same statistically significant threshold of the previous analysis. Only couples with a information of summed FR > 95 percentiles of the bootstrapped distribution, in at least one of the two combinations, were considered.

We then normalized the `Information of Summed FR(E, ISF 12)` generating the information of summed FR index to the sum of the information contained in the mean firing rate of neuron 1 and 2 calculated separately with the following equation:

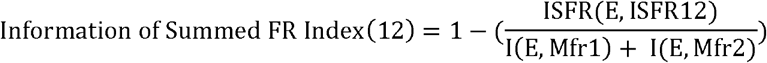

Where `Information of Summed FR Index(12)` is the normalized information carried by the sum of the firing rate of neuron 1 and 2, `ISFR(E, ISFR 12)` and `I(E, Mfrl)` are defined above.

When `Information of Summed FR Index(12)` is equal to 0 it suggests that the information carried by the means of the two neurons are mostly independent, while higher values suggest that the information are more dependent.

### STATISTICAL ANALYSIS

All data are expressed as mean ± standard error of the mean (SEM). Statistical tests were performed using Graphpad Prism 8.0 or SigmaPlot 12.0. Statistical significance was assessed using Wilcoxon Test, Mann-Whitney Test, Z-Test or Chi-square Test, as appropriate. Cumulative distributions were tested using Kolmogorov-Smirnov (K-S) two-sample Test. All statistical analyses were performed on raw data. The level of significance was set at \**p* < 0.05, \*\**p* < 0.01, \*\*\**p* < 0.001.

