## Supplemental figures and tables for "Fast-spiking interneurons of the premotor cortex contribute to action planning"

**Figure S1.**

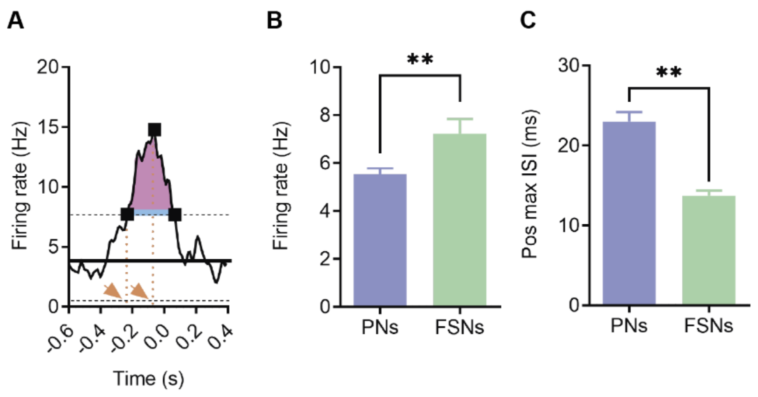


(A) A representative peristimulus time histogram. The black line represents the mean firing rate calculated during resting periods, black dotted lines the upper and lower threshold. The three black squares indicate the first, the maximum and the last point over the threshold. The orange dotted lines and the orange arrows indicate the onset of the activity and the peak time, respectively. The blue line shows the duration of the activity, representing the time over the threshold. The pink area is the area above the threshold. The intensity of activation is defined as the pink area divided by duration of the activity.

(B, C) Mean firing rate (B) and maximum position of interspike intervals (ISI, C) of PNs and FSNs. K-S Test, $^{**}p < 0.01$.

**Figure S2.**

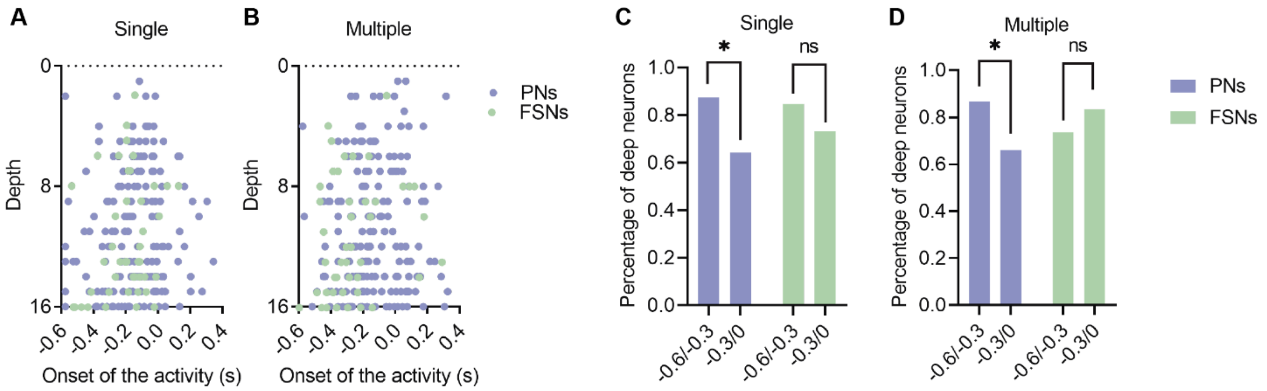


(A, B) PNs (violet) and FSNs (green) depth distribution (across sixteen channels probe) of the onset of the activity in a 1 s window (0.6 s before and 0.4 s after the licking event) during single (A) and multiple (B) licks.
(C, D) Percentage of deep PNs (violet) and FSNs (green) having their onset in two time segments (- 0.6 s to - 0.3 s, and - 0.3 s to 0.0 s) before the licking event, during single (C) and multiple (D) licks. Chi square test, *$p < 0.05$.

**Table S1.**

| | Total Recorded Units | Modulated Units | PNs | FSNs |
|---|---|---|---|---|
| Acute Exp | 693 | 251 | 203 | 48 |
| Chronic Exp | 759 | 373 | 313 | 60 |

Total number of recorded units during acute and chronic experiments. The modulated PNs and FSNs are also reported.

**Table S2.**

| | Lick Enh | Lick Supp | Lick Enh / FP Supp | Lick Enh / FP Enh | Lick Supp / FP Supp | Lick Supp / FP Enh | FP Supp | FP Enh |
|---|---|---|---|---|---|---|---|---|
| PNs | 52 | 31 | 31 | 96 | 55 | 27 | 6 | 15 |
| FSNs | 7 | 2 | 9 | 31 | 11 | - | - | - |

Number of neurons in different functional classes. Lick, licking; FP, forelimb pulling; Enh, enhanced; Supp, suppressed.
